# DSTG: Deconvoluting Spatial Transcriptomics Data through Graph-based Artificial Intelligence

**DOI:** 10.1101/2020.10.20.347195

**Authors:** Jing Su, Qianqian Song

## Abstract

Recent development of spatial transcriptomics (ST) is capable of associating spatial information at different spots in the tissue section with RNA abundance of cells within each spot, which is particularly important to understand tissue cytoarchitectures and functions. However, for such ST data, since a spot is usually larger than an individual cell, gene expressions measured at each spot are from a mixture of cells with heterogenous cell types. Therefore, ST data at each spot needs to be disentangled so as to reveal the cell compositions at that spatial spot. In this study, we propose a novel method, named DSTG, to accurately deconvolute the observed gene expressions at each spot and recover its cell constitutions, thus achieve high-level segmentation and reveal spatial architecture of cellular heterogeneity within tissues. DSTG not only demonstrates superior performance on synthetic spatial data generated from different protocols, but also effectively identifies spatial compositions of cells in mouse cortex layer, hippocampus slice, and pancreatic tumor tissues. In conclusion, DSTG accurately uncovers the cell states and subpopulations based on spatial localization.

## Introduction

Cells of different types are spatially and structurally organized within tissues to perform their functions. Uncovering the complex spatial architecture of heterogenous tissue is significant for understanding the cellular mechanisms and functions in diseases. The fast advance of single-cell RNA sequencing technologies (scRNA-seq) attracts the attention to elucidate the heterogenous cell formation [1–4] and trace the lineage relationship within tissue [5–7]. Unfortunately, due to the lack of spatial information, scRNA-seq is incapable of identifying the structural organization of heterogeneous cells within a complex tissue. Therefore, as the complementary to scRNA-seq, spatially resolved transcriptomic profiling methods [8–10] have been introduced. To reveal the spatial cytoarchitectures within tissues, sequencing-based high-throughput spatial transcriptomics (ST) technologies [11–14], such as 10x Genomics Visium [8] and Slide-seq [15, 16], use spatially indexed barcodes with RNA sequencing that allows quantitative analysis of the transcriptome with spatial resolution in individual tissue sections.

Emerging ST technologies are able to spatially index transcripts and measure expression profiles, advancing our understanding of precise tissue architectures. However, the resolution of ST data is far lower than single cell level. Transcripts captured at a specific location by a “spot”[8] or a “bead”[15, 16] is usually composed of a mixture of heterogenous cells. For example, Visium, one of the microarray-based spatial transcriptomics techniques developed by 10X Genomics, uses spots of 50 μm diameter, with each spot covering 10-20 cells in average, which varies depending on the tissue histology [17]. Even for the Slide-seq [15, 16] that quantifies gene expression with high resolution (10 microns), one pixel may still be overlapped with multiple cells. As a result, the measured gene expressions at a “spot” reflects a mixture of cells. Therefore, uncovering the cell compositions within each spot of the spatial transcriptomics data is critical for investigating tissue’s molecular and cellular architecture at high resolution.

To address this problem, very few tailored approaches have been developed yet. SPOTlight [18] is a deconvolution algorithm using non-negative matrix factorization regression and nonnegative least squares, which has been applied to ST data [16] successfully. Specifically, SPOTlight incorporates the reference scRNA-seq data to identify cell type-specific topic profiles, which is further used to deconvolute spatial spots. This method leverages scRNA-seq data for the identification of cell states and subpopulations to deconvolute the spatial transcriptomics data, showing that leveraging well-characterized scRNA-seq data will aid and facilitate the exploration of spatial datasets. A major limit of this ST deconvolution method is that the intrinsic topological information of cell type constitutions within spots, which provides crucial information about the relations between the observed gene expression patterns and associated cell types at spots, cannot be effectively learned and utilized.

In recent years, graph convolutional networks (GCN) [19] has demonstrated promising capability in utilizing such intrinsic topological information of data to improve model performance. The topological relations inside the data, such as similarity between samples, can be represented as graphs. Through learning the shared kernel used in spectral graph convolution across all nodes in a graph, a semi-supervised GCN model captures local graph structures as well as node features and incorporates both information as latent space representation. Graph Convolutional Networks (GCN) [19] and its variants [20, 21] have been applied to different scenarios successfully, including cancer patient subtyping using real-world evidence [22], protein prediction [33] and drug design [23], as well as single cells and diseases [24–28]. These work shows that, through effectively learning and leveraging the latent representation and topological relations among data, GCN models are able to significantly improve model performance.

In this work, we have developed a novel graph-based artificial intelligence model, Deconvoluting Spatial Transcriptomics data through Graph-based convolutional networks (DSTG), for reliable and accurate decomposition of cell mixtures in the spatially resolved transcriptomics data. Based on the well-characterized scRNA-seq dataset, DSTG is able to learn the precise composition of spatial transcriptomics data using semi-supervised graph convolutional network. The performance of DSTG has been validated on synthetic ST data, as well as on different experimental ST datasets with well-defined structures including mouse cortex layer, hippocampus tissue, and pancreatic tumor tissues. In addition, we provide the implementation software of DSTG as a ready-to-use Python package, which is compatible with current spatial transcriptomics profiling datasets for accurate cell type deconvolution.

## Results

### Overview of DSTG

Herein, we propose a novel, graph-based artificial intelligence approach, namely DSTG, to deconvolute spatial transcriptomics data through graph-based convolutional networks. The DSTG approach leverages scRNA-seq data to unveil the cell mixtures in the spatial transcriptomics data (**Figure 1**). Our hypothesis is that the captured gene expression on a spot is contributed by a mixture of cells located on that spot. Our strategy is to use the scRNA-seq-derived synthetic spatial transcriptomics data called “pseudo-ST”, to predict cell compositions in real-ST data through semi-supervised learning. First, DSTG constructs the synthetic pseudo-ST data from scRNA-seq data as the learning basis of our method. Then DSTG learns a link graph of spot mapping across the pseudo-ST data and real-ST data using shared nearest neighbors. The link graph captures the intrinsic topological similarity between spots and incorporate the pseudo-ST and real-ST data into the same graph for learning. Then, based on the link graph, semisupervised GCN is used to learn a latent representation of both local graph structure and gene expression patterns that can explain the various cell compositions at spots. The major advantages of such similarity-based semi-supervised GCN model are: 1) sensitive and efficient, since for each spot, only the features of similar spots (i.e., neighbor nodes) are used; and 2) acquiring generalizable knowledge about the association between gene expression patterns and cell compositions across spots in both pseudo- and real-ST, since the weight parameters in the convolution kernel are shared by all spots. To test the performance of DSTG, we use synthetic pseudo-ST data generated with cell mixtures of known cell type compositions from peripheral blood mononuclear cell (PBMC) scRNA-seq datasets [29], in which DSTG presents superior performance than the SPOTlight method. Furthermore, DSTG is validated and applied to real tissue context from mouse cortex, hippocampus, and pancreatic tissues with well-defined structures.

**Figure 1.**
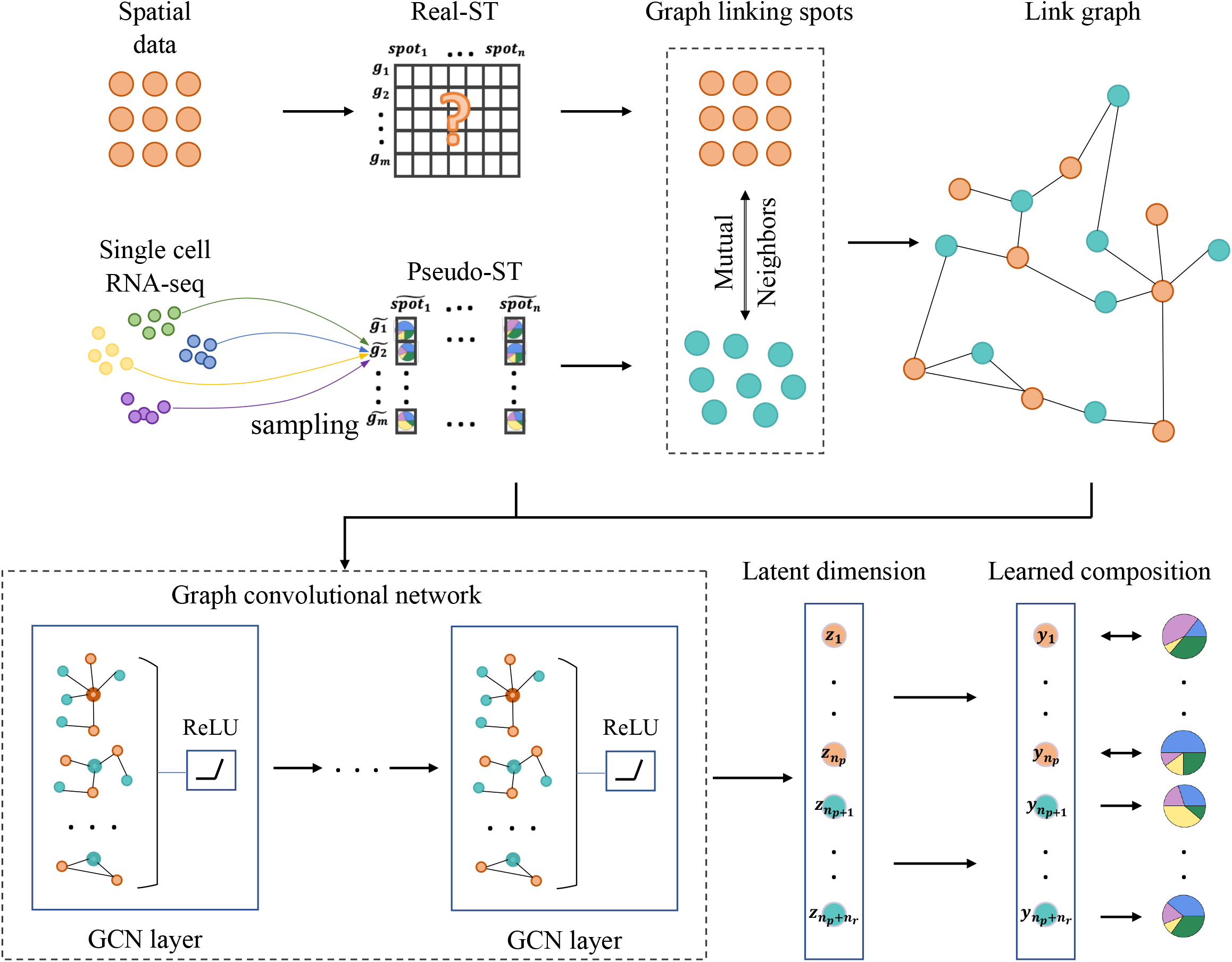
Schematic overview of DSTG for deconvoluting spatial transcriptomics data. Schematic representation of how DSTG deconvolutes spatial transcriptomics (ST) data using scRNA-seq profiles. DSTG first generated a pseudo-ST data with cell mixtures from scRNA-seq data. Between this pseudo- and real-ST data, DSTG identifies a link graph of spot mapping. Based on the link graph, graph convolutional network is used to propagate both pseudo-ST and real-ST data into the latent layer and identify the portions of different cell types for each spot. In this way, cell compositions of real-ST data can be predicted and learned from pseudo-ST data.

### Performance on benchmarking data

To evaluate the performance of DSTG, we use the synthetic spatial data generated by scRNA-seq cell mixtures as ground truth. Briefly, each spot of this synthetic spatial data is constructed by combining the randomly selected 2 to 8 cells from scRNA-seq data. Such synthetic spatial data not only mimics real spatial transcriptomics data, but also provides ground truth that can be used to evaluate the DSTG’s performance in identifying the proportions of different cell types within each synthetic spot. As for the evaluation metrics, we use the Jensen-Shannon Divergence (JSD), which is a distance metric that measures the similarity between two probability distributions. A smaller value of JSD represents a higher similarity between two distributions, thus signifies a higher accuracy of estimated cell type compositions across spots.

Specifically, we use 13 PBMC scRNA-seq datasets [29] profiled by different protocols, with well-characterized cell populations and discrete cell numbers, to generate benchmarking synthetic spatial data. For each PBMC data, we generate 10 synthetic data and apply both DSTG and SPOTlight to those 10 synthetic data for comparison. Our results show that DSTG achieves lower JSD values (mean JSD = 0.12, **Figure 2A**), which is significantly lower (*P* value < 2.2e-16) than SPOTlight (mean JSD = 0.24), indicating the higher accuracy of DSTG than SPOTlight across datasets generated from different experiment protocols. Notably, DSTG shows the most accuracy than SPOTlight in the CEL-Seq2 synthetic datasets. Though SPOTlight performs the best on Quartz-Seq2 datasets, DSTG still outperforms SPOTlight with lower JSD value. In addition to PBMC, to examine the performance of DSTG on other different tissues, we include 8 other scRNA-seq data from different tissues and protocols to generate the benchmarking synthetic data. Then we compare DSTG with SPOTlight based on the synthetic data from these 8 additional scRNA-seq data. As shown in **Figure 2B**, the predicted results of DSTG still outperform SPOTlight using the JSD evaluation metric. Notably, DSTG achieves the mean JSD values of 0.016 and 0.087 for the Smart-seq2 and snRNA-seq datasets, respectively, which are better than the ones achieved by SPOTlight (0.19 and 0.24). These consistently superior performances of DSTG demonstrate the accuracy and robustness of our method.

**Figure 2.**
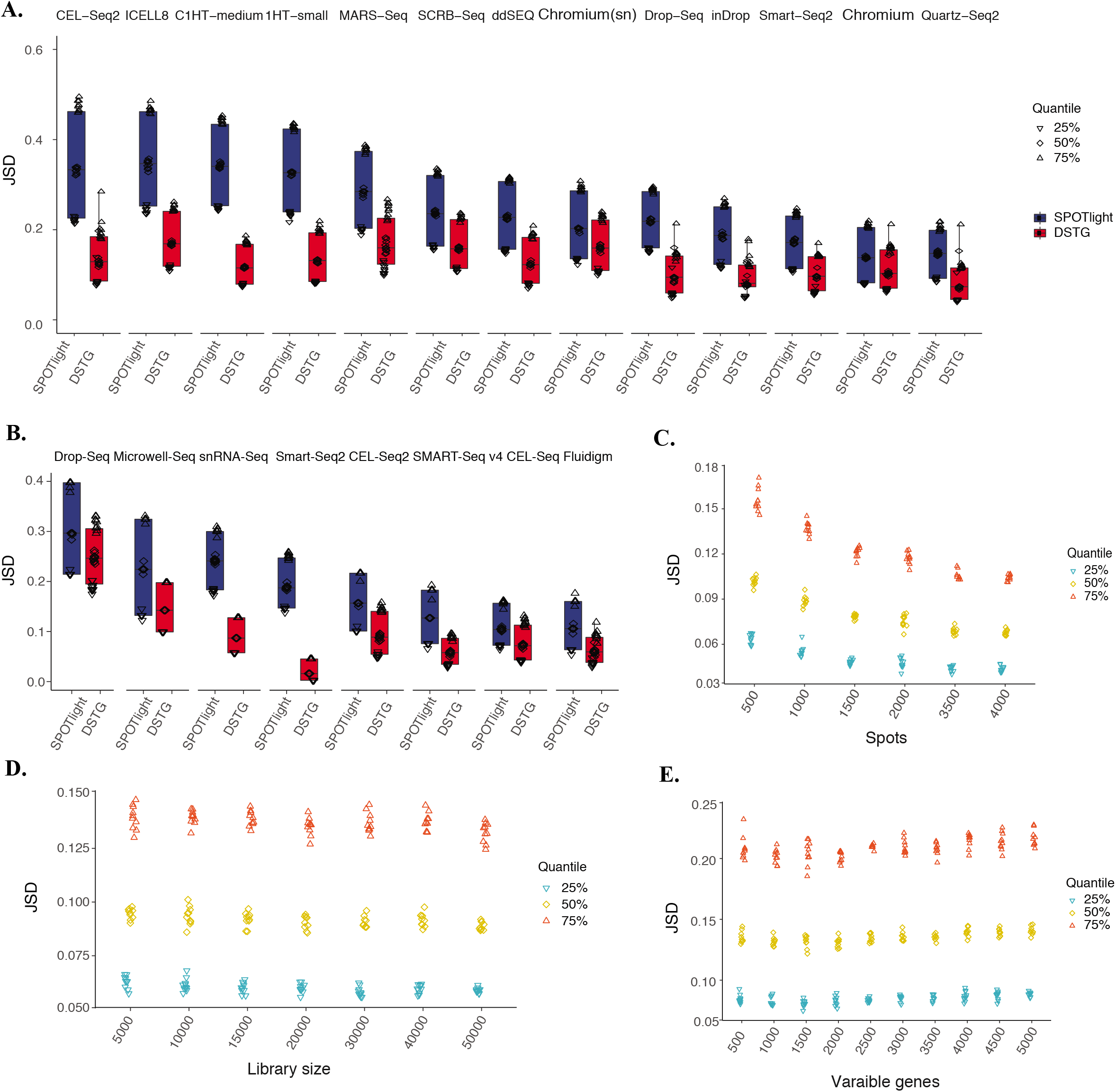
Performance of DSTG on benchmarking datasets. **A.** Performance of DSTG is assessed and compared with SPOTlight by synthetic spatial data generated from 13 PBMC datasets of different scRNA-seq protocols. **B.** Performance of DSTG is further benchmarked with SPOTlight by synthetic spatial transcriptomics data generated from the other scRNA-seq datasets from different tissues and protocols. **C.** DSTG’s performance on synthetic data with different number of spots. **D.** DSTG’s performance on synthetic data with different library depths. **E.** DSTG’s performance on synthetic data with different number of variable genes. In **A**-**E**, the y-axis represents the JSD value.

To investigate whether DSTG is sensitive to the design of synthetic data, we generate discrete synthetic data with different number of spots, library sizes, and variable genes, which covers the characteristics of the current and emerging ST data. For the synthetic data with different spot numbers (500 - 4,000) (**Figure 2C**), we find that DSTG tends to perform better with more spots in the synthetic data, suggesting that the more spots used, the better the model is trained. Meanwhile, the result suggests that using 1,500 spots is sufficient to reach high performance in practice, as the marginal gain of performance is neglectable when using more spots. For the synthetic data with different downsampled library sizes (5,000 - 50,000 reads per cell) (**Figure 2D**), DSTG shows stable accuracy at lower or higher library sizes. For the synthetic data with different number of variable genes (500 - 5,000) (**Figure 2E**), DSTG demonstrates stable performance, with optimal performance reaches at 2,000 variable genes.

### Spatial decomposition of mouse cortex layer

To examine whether DSTG can reveal microanatomical structures in complex tissue, we use the 10X Visium ST data of cerebral cortex layer in mouse brain. This cortex layer has well-defined cytoarchitecture and thus is suitable to evaluate DSTG’s performance. To deconvolute this ST data by DSTG, we use the scRNA-seq dataset profiled by the SMART-Seq2 protocol from the Allen Institute, which consists of ~14,000 adult mouse cortical cell taxonomy and 22 cell types (**Figure 3A**). With this scRNA-seq data, the spatial deconvolution of the ST data by DSTG accurately reconstructs the architecture of brain cortex layer (**Figure 3B**). The identified heterogenous cell proportions of each localized spot are shown by the pie chart at the respective spot, which is confirmed by their existence in cortical areas, suggesting the high accuracy and sensitivity of the DSTG’s predictions. Moreover, our predicted compositions provide more detailed information about the heterogeneity of this area. Specific investigation shows the regional enrichment of each cell type based on their identified proportions. Illustrative examples are the differentially enriched neuronal subtypes including cortical layer 2/3 (L2/3), cortical layer 6 (L6b), and oligodendrocytes (oligo) (**Figure 3C**). The subpopulation of L2/3 are shown with high compositions in the outside liner of spots within the cortex. Spots with most L6b cells are shown with high proportions in the inner liner of the cortex. Towards the innermost layer, oligodendrocytes cells are mainly abundant in these spots. These data are consistent with the layered cytoarchitecture of the cortex tissue. The ability to identify the distinct spatial cellular compositions of each spot in the cortical neuronal layers indicates the accuracy and sensitivity of DSTG.

**Figure 3.**
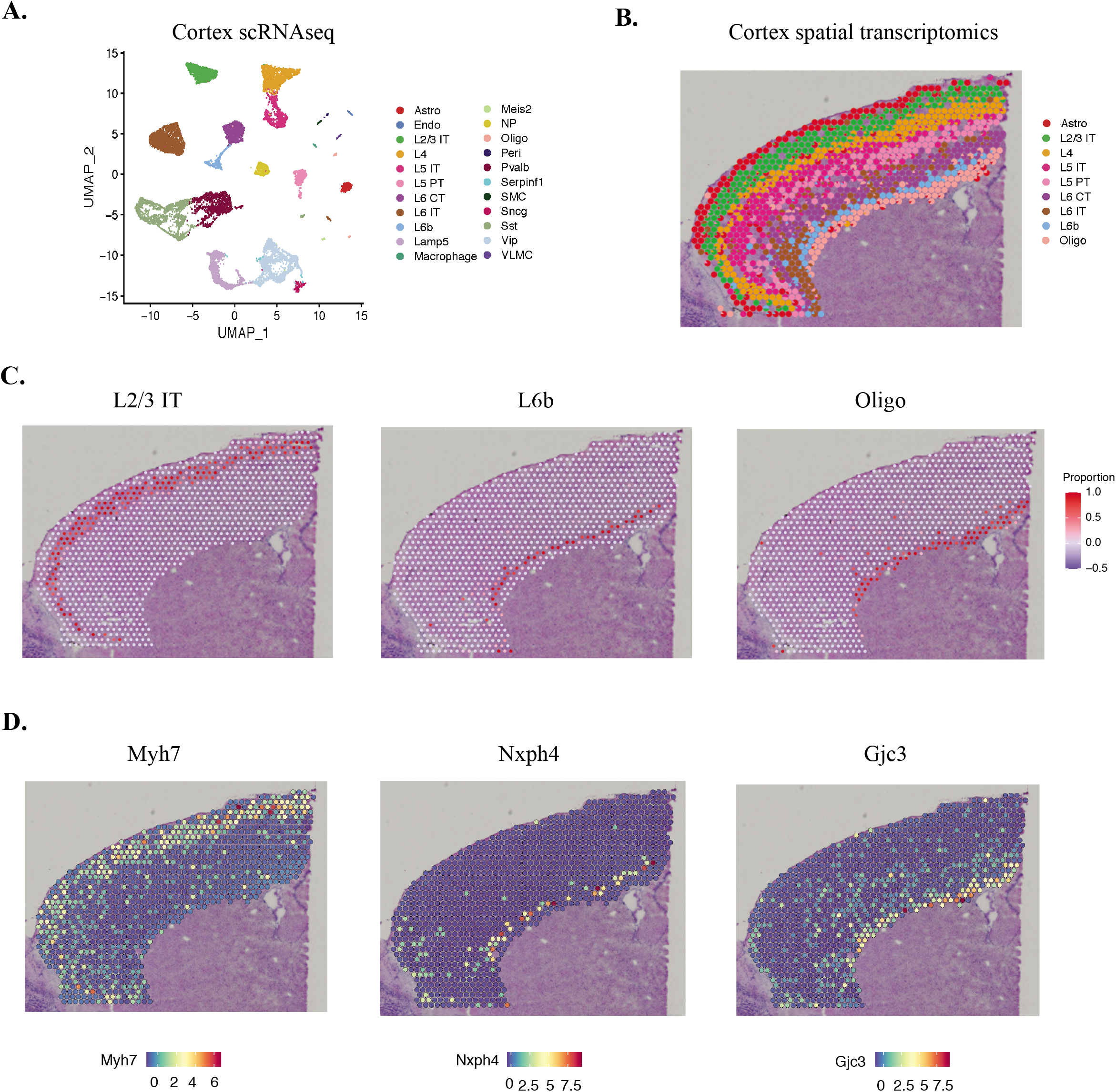
Spot deconvolution of mouse cortex layer. **A.** UMAP projections of single cell RNA-seq data from mouse cortex tissue. Different cell types are labeled and colored according to known cell annotations. **B.** Spatial plot with pie chart shows the predicted cell compositions of each spot within the cortex layer. **C.** Spatial plot shows the proportions of specific neuron subtypes in the spots within the captured region. Red spot indicates high proportion of the respective cell type. **D.** Visualization of the spatial expressions of cell type specific markers in ST data. Red color indicates the high gene expression in that spot.

To examine if the cell type specific genes are enriched in their corresponding spatial locations, we investigate the distribution of marker genes known to be specific to the respective cell types (**Figure 3D**). For example, the top expressed gene marker of L2/3 cells, Myh7, is detected with high expression in the ST data of L2/3 dominated spots, which is in line with the predicted proportions of this cell type. The top markers of L6b and Oligo, Nxph4 and Gjc3, also show high expression in their corresponding spots respectively. Meanwhile, these genes are undetectable or detected at very low levels at other spatial spots. It’s worth noting that the partial expression of cell type markers in a specific ST spot may reflect the heterogeneous composition of cell types in that spot. Interrogation of these differential genes further confirms the accurate predicted cell proportions within spots in the tissue section.

### Mapping distinct cell populations of mouse hippocampus

The fast advance of spatial transcriptomics technologies raises new challenges in deconvoluting the transcriptomics data at each spot: spots become smaller, meanwhile the total number of spots grows exponentially, but the sequencing depth at each spot becomes much lower. We demonstrate the performance of DSTG on such emerging ST data, using the recently available Slide-seq v2 [15] of mouse hippocampus tissues as an example. Comparing with the 10x Genomics’ Visium platform, the bead size of the Slide-seq v2 platform is 5.5-fold smaller, thus the spatial resolution is 25 ~ 100-fold higher. Consequently, the typical median library size per bead is 550 UMIs (Unique Molecular Identifiers), 100-fold lower than 10X Genomics Visium.

To deconvolute this ST data, we use the existing single-cell RNA-seq dataset from mouse hippocampus [30], which consists of 52, 846 cells with 19 cell types that are profiled by the Drop-seq protocol (**Figure 4A**). Based on this scRNA-seq data, DSTG’s spatial decomposition of the ST data accurately identifies different cell types within the hippocampus slice (**Figure 4B**). The spatially localized pie charts represent the identified different cell proportions in the slice. Moreover, our predicted compositions provide more detailed information about the heterogeneity of this area. Closer investigation confirms the regional enrichment of specific cell types with their identified composition (**Figure 4C**). For example, oligodendrocyte cells are identified with high compositions in the middle wide strips within the hippocampus slice. In the Cornu Ammonis (CA) subfield, CA3 principal cells scatter within the slice with low proportions, but majorly abundant in the half strip at the right of the slice. Ependymal cells present mainly at an irregular circle and the other band at the top right of the slice, which data are consistent with the spatial structures within the mouse hippocampus.

**Figure 4.**
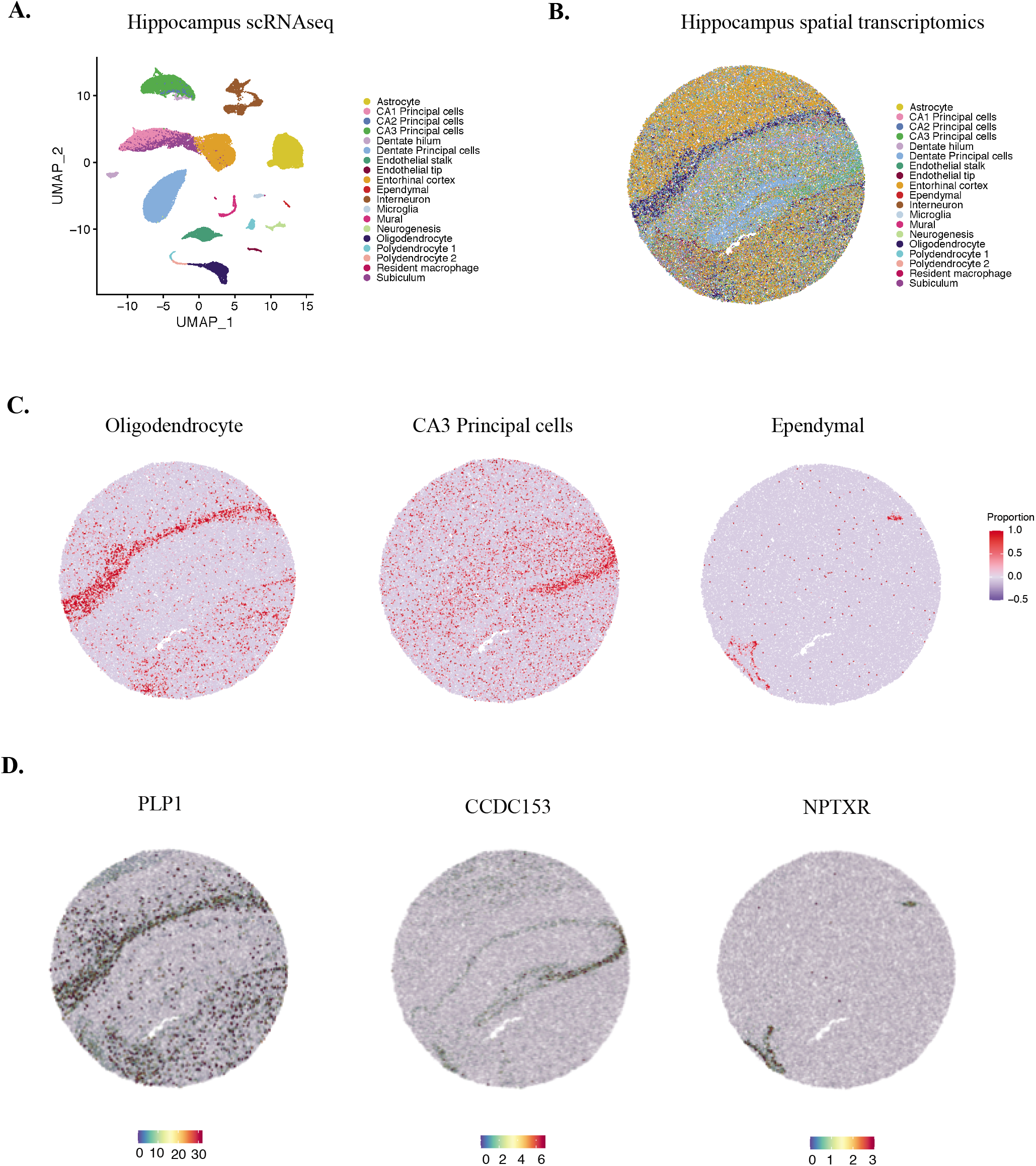
Spatial chart of mouse hippocampus tissue using DSTG. **A.** UMAP projections of single cell RNA-seq data from mouse hippocampus tissue. Different cell types are labeled and colored based on known cell annotations. **B.** Spatial plot with pie chart shows the predicted cell compositions within the captured locations in the mouse hippocampus. **C.** Spatial plot presents the proportions of specific neuron subtypes within the captured location. Red color indicates high abundance of certain cell type in this region. **D.** Spatial expression of cell type specific markers of the respective neuron subtypes in the ST data.

To further evaluate our method, we select the cell type specific genes from scRNA-seq data and assess their expression in the ST data. As expected, the top differentially expressed gene marker in oligodendrocyte (**Figure 4D**), PLP1, expresses strictly in line with the oligodendrocyte region based on the ST data. CCDC153 is the top expressed gene marker in CA3 principal cells that is also detected with high expression in the ST data of CA3 principal cells. Another example is NPTXR, the marker of ependymal, is enriched at regions with a high abundance of Ependymal cells. These cell specific genes detected in their corresponding locations further underlines the accuracy of DSTG. In summary, DSTG demonstrates accurate and reliable deconvolution capabilities on ST data generated from the latest ST platform such as the Slide-seqV2, which has much smaller spot size, much larger number of spots, and much lower sequencing depth on each spot.

### Deconvolution of pancreatic cancer tissue sections

Tissues in diseases such as tumors exhibit unique pathological cytoarchitectures. To further demonstrate and test the DSTG’s performance in such conditions, we apply it to two ST data obtained from two tumor sections of pancreatic ductal adenocarcinoma (PDAC), i.e. PDAC-A and PDAC-B. Sample-matched scRNA-seq data (**Figure S1**) generated by inDrop protocol is used by DSTG to deconvolute the respective ST data in samples PDAC-A and PDAC-B respectively.

For the PDAC-A sample (**Figure 5A**), after DSTG’s deconvolution, we observe discrete regional enrichment of cancer clones and non-cancer cells. Specifically, cells of cancer clone S100A4 and TM4SF1 are mainly identified mixed in the spots of cancerous region, which are excluded from the spots of ductal cells including the centroacinar ductal cells and the co-localized antigen presenting ductal cells. Stroma cells are involved between the ductal cells and cancer cells, which are consistent with previous results annotated by hematoxylin and eosin (H&E) staining and brightfield imaging [13]. We also find a few proportions of hypoxic ductal cells in the spots close to the cancerous region, indicating the low oxygen environment in tumor. Further inspections of specific cell types (**Figure 5B**) including cancer clone cells and ductal cells, confirm their regional proportions on their identified structures. The point size and related color indicate different compositions in the spatial spots.

**Figure 5.**
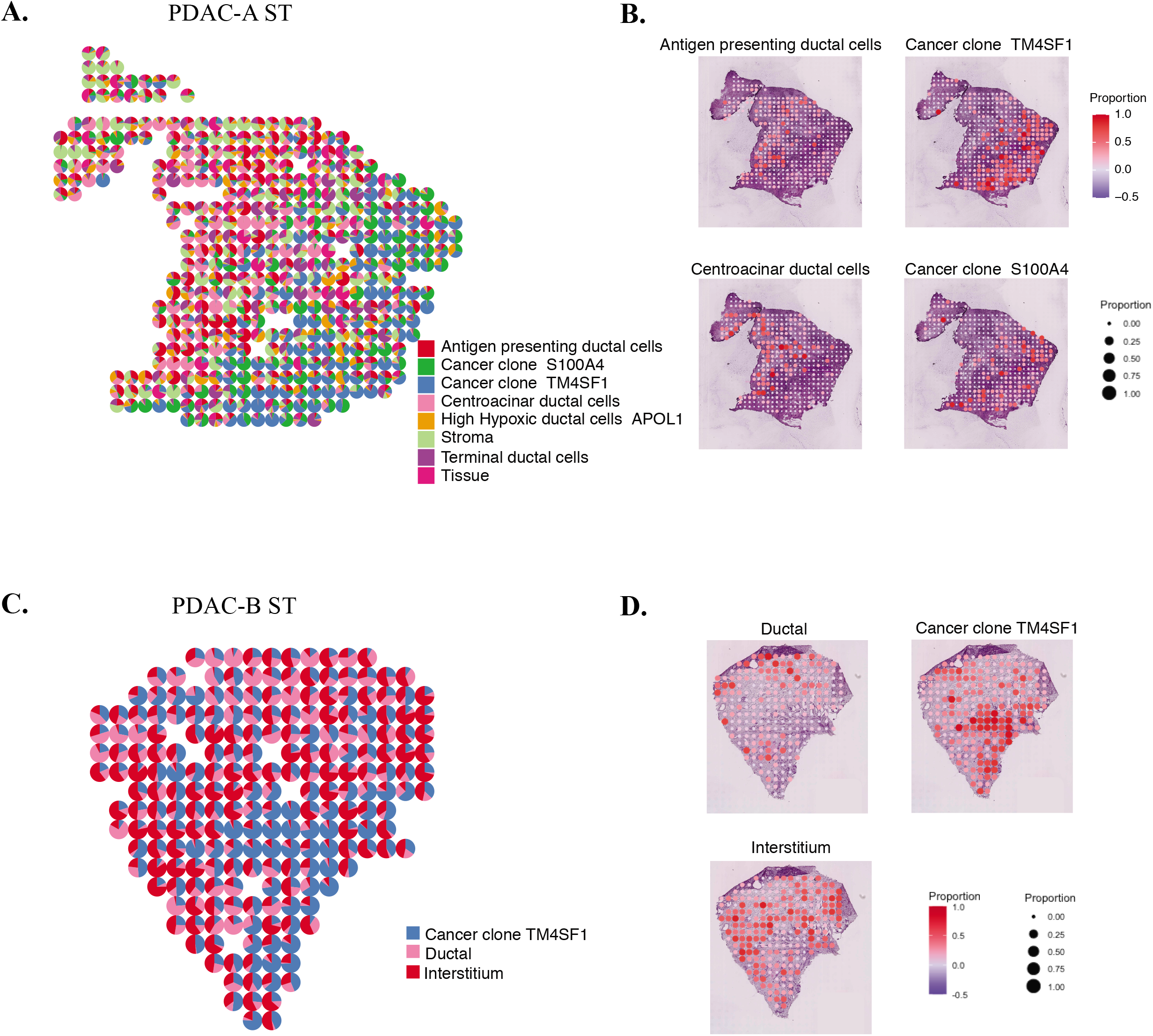
Mapping spatial spots across pancreatic cancer tissue. **A.** Spatial plot shows the predicted compositions of different cell types within the captured spots of PDAC-A tissue slice. **B.** Predicted proportions of different cell types including antigen presenting ductal cells, centroacinar ductal cells, cancer clone TMSF1, and S100A4. **C.** Spatial plot presents the predicted compositions of different cell types within captured spots of PDAC-B tissue slice. **D.** Predicted proportions of different cell types including ductal cells, cancer clone TMSF1, and interstitium. Red color indicates high proportion of certain cell type.

In the other PDAC-B sample, as shown in **Figure 5C**, cells of cancer clone TM4SF1 rather than cancer clone S100A4 are identified. These cancer clone TM4SF1 cells are localized preferentially in the spots of the bottom right region, distinguished from the interstitium and ductal cells. We notice that most interstitium cells are adjacent to the cancer clone TM4SF1 cells, and some interstitium cells co-localize with the cancer cells. Ductal cells are mainly abundant in the spatial spots of the top region. These findings highlight the precise consistency with previous H&E staining results [13]. Further inspections of cancer clones and ductal cells (**Figure 5D**) confirm their regional compositions on their known locations. These results of DSTG are consistent with independent histological annotations, supporting its ability to identify accurate cellular compositions from the ST data of tumor tissues.

In conclusion, DSTG is able to detect the unique cytoarchitectures in diseased tissues, distinguishes the spatial distribution of tumor cells evolved from different clones, and characterizes cancer-specific cellular phenomena such as local hypoxia as well as the suppression of antigen presenting in the tumor cell dominating regions.

## Discussion

Spatial transcriptomics provides unprecedented opportunities to study tissue heterogeneity and cell spatial organization [31–33]. However, the resolution of spatial transcriptomics is less than the single-cell level. As single spot in spatial transcriptomics data may cover heterogenous cell types, our DSTG method aims to determine the proportions of different cell types and states across spots where genes are reliably identified. In this study, we present the DSTG method for performing cell type deconvolution in spatial transcriptomics data using the graph convolutional networks. DSTG is evaluated by benchmarking synthetic data generated from PBMC and other tissues, in which DSTG demonstrates excellent accuracy between the predicted cell mixtures and the actual cell composition. DSTG is also shown to achieve high consistency with H&E staining observations on spatial transcriptomics data from complex tissues including mouse cortex, hippocampus and human pancreatic tumor slices.

As SPOTlight is also used to deconvolute the spatial transcriptomics data, we compare DSTG with SPOTlight, and find DSTG consistently outperforms SPOTlight on benchmarking synthetic data. From a technical perspective, DSTG provides some major advantages. First, DSTG simultaneously utilizes variable genes and graphical structures through a non-linear propagation in each layer, which is appropriate for learning the cellular composition due to the heteroskedastic and discrete nature of spatial transcriptomics data. Second, DSTG identifies the respective weights of different cell types in the pseudo-ST data generated from scRNA-seq data, which can be effectively leveraged to learn the cell compositions in the real-ST transcriptomics data. Third, as the sequencing depth of spatial data is expected to increase, DSTG has been shown to perform better to interrogate the cell distribution quantitively in such spatial transcriptomics data.

In addition to the successful results, there are several aspects that DSTG can be improved. First, as an artificial intelligence (AI) model, DSTG shows not only the merits of its kind, but also some limitations including the black-box nature of AI models [34–36], which can be addressed through downstream analysis that can ameliorate some of the problems and bring insights into the learned cellular compositions. Second, as a graph model, improving the built graph can further boost the model performance. Our link graph based on mutual nearest neighbors best captures the spots’ similarity in spatial data, reflecting the effective graph representation. As a fast-growing research field, new approaches of building graph are emerging, of which we will test and adapt in future versions of DSTG.

Our DSTG method paves the way for inferring functional relationships between heterogenous cell subpopulations based on their composition and colocalization in the tissue spots. This includes intercellular communication across neighboring spots, which opens up future possibilities of studying the complete interactome in a spatially resolved manner. Moreover, as the precise composition of tissue may vary from one individual patient to the other, the spatial composition of cellular subpopulations can be of prognostic value for patients in the future. We anticipate that the spatial deconvolution using DSTG will contribute to future patient prognosis and pathological assessments. Overall, DSTG demonstrates as a robust and accurate tool to determine cell-type locations and precise compositions of spatial spots, which provides an unbiased perspective and investigation into the spatial organization of distinct cellular populations in tissue.

## Methods

### Variable gene selection

For the scRNA-seq data, we first identify genes that exhibit the most variability across different cell types using the analysis of variance (ANOVA). The top 2,000 most variable gene features in the scRNA-seq data are selected according to adjusted *P* values with Bonferroni correction. Using the scRNA-seq data of the top variable genes, we then generate the pseudo-ST data with synthetic mixtures of cells with known cell compositions. The gene expressions at each pseudospot of the pseudo-ST data is generated by combining the randomly selected 2 to 8 cells from the scRNA-seq data. The library size is limited to 20,000 UMI counts for the pseudo-ST data. For simplicity and illustration, we consistently use the term “spot” to represent the synthetic cell mixture of the pseudo-ST data as well as a spot or a bead of real-ST data.

### Link graph

For both pseudo-ST data and real-ST data, we first perform the standardized transformation, *i.e.*

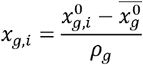

where 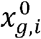 is the raw counts of gene *g* and spot *i*, 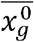 is the mean of 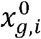 over all spots, and *ρ_g_* is the standard deviation of 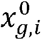. Thus *x_g,i_* is the standardized gene expression.

After the data standardization, we then build a link graph incorporating pseudo-ST and real-ST data for the DSTG method. The built graph is *G* = (*V,E*) with *N* = |*V*| nodes denoting the spatial spots and *E* representing the edges. *A* is the adjacent matrix in terms of this graph. Here we apply the dimension reduction of the pseudo-ST and real-ST data by canonical correlation analysis, and then identify the mutual near neighbors [37] in the space of reduced dimension.

First, with the pseudo-ST data and the real-ST data represented as *X_pseudo_^m×n_p_^* and *X_real_^m×n_r_^* where *m* is the number of variable genes, and *n_p_* and *n_r_* are the respective number of spots, we project these two data into a lower *S* dimension space by canonical correlation vectors *μ_s_* of *n_p_* dimension and *v_s_* of *n_r_* dimension, where *s* = 1, ⋯, *S*, to maximize

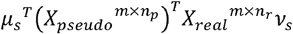
subjecting to the constraints 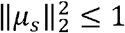 and 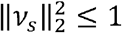. To identify the canonical correlation vector pairs, we use singular value decomposition and get the S’ canonical correlation vector pairs with the S’ largest eigenvalues. Each pair of *μ_s_* and *v_s_* projects the original data *X_pseudo_^m×n_p_^* and *X_real_^m×n_r_^* to the *s*-th dimension of the low dimension space. For DSTG, we take *S* as 20 for the reduced dimension space.

Second, in the low dimension space, we identify the mutual nearest neighbors among spots from pseudo-ST and real-ST data. Specifically, if spot *i* is in one of nearest neighbor of spot *j* by *k*-nearest neighbor (KNN, default *k* is 200), meanwhile spot *j* is in one of nearest neighbor of spot *i* by KNN, then spot *i* and spot *j* are mutual nearest neighbors. In this way, we build the link graph between the pseudo-ST data and the real-ST data. To further utilize the information of real-ST data in the DSTG model, we also identify the mutual nearest neighbors within the real-ST data itself. To this end, the final link graph is built and represented by the adjacent matrix *A*. That is, if spot *i* and the other spot *j* are mutual nearest neighbors, *A_ij_* = 1, otherwise *A_ij_* = 0. This graph captures the intrinsic topological structure of spot similarity between all spots.

### DSTG method

We utilize the graph convolutional network on the link graph *G* = (*V,E*) for the identification and prediction of the compositions of different cell types in the spatial transcriptomics data. Each spot is viewed as a node. The cell mixtures in the pseudo-ST data are generated with known compositions. The goal of DSTG is to predict the cell type compositions of the real spatial data by using not only the features of each spot but also the graph information leveraging the pseudo-ST data and real-ST data, which are characterized as the above adjacent matrix *A*. Explicitly, the DSTG method takes two inputs. One input is the spot similarity graph structure learned above (see the **Link graph**). The other is the data matrix of combined pseudo-ST and real-ST data. As denoted above, with the pseudo-ST data and the real-ST data represented as *X_pseudo_^m×n_p_^* and *X_real_^m×n_r_^* where *m* is the number of variable genes, while *n_p_* and *n_r_* are the corresponding number of spots, the input data matrix is shown as

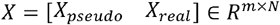
where *N* – *n_p_* + *n_r_*.

Herein, with these two inputs, i.e. *X* and *A* the DSTG is constructed with multiple convolutional layers. For efficient training of DSTG, the adjacent matrix *A* is modified and normalized as:

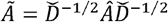
where *Â* – *A* + *I,I* is the identity matrix, and 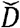 is the diagonal degree matrix of *Â* Specifically, each GCN layer is defined as:

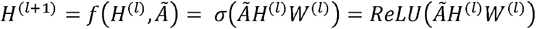
where *H*^(*I*)^ is the input from the previous layer, *W*^(*l*)^ is the weight matrix of the *l*-th layer, *σ*(·) = *ReLU*(·) is the non-linear activation function, and the input layer *H*^(0)^ = *X*. The composition of a specific cell type *f* at a pseudo-spot *i* is represented as *y_i,f_* ∈ *Y_p_*, where *i* ∈ [l,⋯,*n_p_*] and cell type *f* ∈ {1, ⋯, *F*], *F* represents the total number of different cell types, and *Y_p_* ∈ *R^n^_p_^×F^* represents the known cell compositions at all spots from the pseudo-ST data.

Specifically, for a three-layer DSTG with *F* distinct cell types, the forward propagation is realized as:

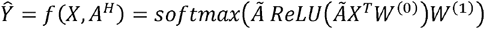
where *W*^(0)^ ∈ *R^m×h^* is the input-to-hidden weight matrix projecting the input data with *m* variable genes into an *h* dimension hidden layer, *ReL,U* stands for the rectified linear unit activation function, *W*^(1)^ *R^h×F^* is a hidden-to-output weight matrix, and 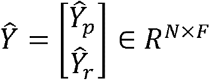 is composed of two components: 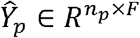 represents the predicted proportions of different cell types at pseudo-spots and 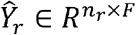 represents the prediction of cell compositions at real-ST spots. The *softmax* activation function below are used as the activation function in the output layer that learns the cell type proportions,

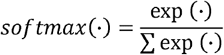

. The evaluation function is defined as the cross-entropy at pseudo-spots, *i.e.*

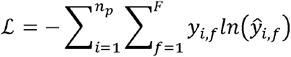
where *y_i,f_* ∈ *Y_p_* and 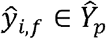. The goal of this semi-supervised learning is to minimize the cross-entropy 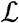 between the known cell compositions *Y_p_* and the predicted cell proportions 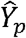. During the propagation of each layer, the model will reduce the cross-entropy error on the training data. After training, we have

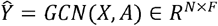

. Note that 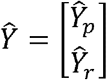. Thus, cell compositions at real spots are predicted as 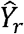.

When applying the DSTG model, we randomly split the pseudo-ST data as training (80%), test (10%), and validation set (10%), while the real-ST data is unlabeled and will be predicted. For the spatial transcriptomics data in this study, we train three-layer DSTG models for a maximum of 200 epochs using the Adaptive Moment Estimation (Adam) algorithm [38] with a learning rate of 0.01 and early stopping with a window size of 10. For the dimension of the latent layer, we screen options of 32, 64, 128, 256, 528, and 1,024 dimensions and select the optimal one.

For the evaluation metrics, we use the Jensen–Shannon divergence (JSD) score, which is a symmetrized and smoothed version of the Kullback–Leibler divergence. With discrete probability distributions *P* and *Q* defined on the same probability space *χ*, the JSD score is defined by

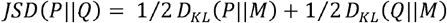
where *M* – 1/2 (*P* + *Q*), and *D_KL_*(*P*||*Q*) represents the Kullback-Leibler divergence from *Q* to *P,* i.e.,

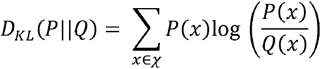

## Data availability

All single-cell RNA-seq datasets are downloaded from their public accessions. The first benchmarking PBMC scRNA-seq datasets [29] in **Figure 2A** are generated by 13 different protocols, including C1HT-medium, ICELL8, MARS-Seq, Chromium (sn), CEL-Seq2, gmcSCRB-Seq, C1HT-small, inDrop, Drop-Seq, ddSEQ, Smart-Seq2, Chromium, Quartz-Seq2. These 13 PBMC datasets are publicly available through the Gene Expression Omnibus (GSE133549). To evaluate the impact of synthetic data (**Figure 2C-E**), we use the Smart-Seq2 PBMC dataset to generate discrete synthetic data with different number of spots and variable genes, as well as different sequencing depths.

For the second benchmarking in **Figure 2B**, the Drop-Seq data is downloaded from the Short Read Archive under accession number SRP073767 [2]. The Microwell-Seq data is profiled using the Microwell-Seq protocol [39] that can be downloaded from the Mouse Cell Atlas. The snRNA-Seq data is profiled from the entorhinal cortex from human brains of Alzheimer’s Disease, yielding a total of 13,214 high quality nuclei [40] using the single-nucleus RNA-seq protocol, which can be downloaded from GEO (accession number GSE138852). The Smart-Seq2 data is profiled using the Smart-Seq2 platform [41], which is profiled from melanoma tumor and downloaded from GEO (accession number GSE72056). The CEL-Seq2 data is obtained from human cadaveric pancreata using the CEL-Seq2 protocol (accession number: GSE85241) that consists of 2,122 cells and 18,915 genes [42]. The SMART-Seq v4 data is downloaded from dbGAP (accession number phs001790) [43], which is generated using SMART-Seq v4 platform. This dataset contains 16024 genes and 14055 cells, from 34 cell types in the middle temporal gyrus of human cerebral cortex. The CEL-Seq data is obtained from 3 human lung adenocarcinoma cell lines using the CEL-Seq platform that consists of 570 cells and 12,627 genes [44], which can be downloaded from GEO (accession number: GSE117617). The Fluidigm data is profiled using Fluidigm C1 platform with 11,778 cells and 3,803 genes [45]. We download this data from GEO (accession number: GSE81608).

## Acknowledgements

The authors acknowledge the DEMON high performance computing (HPC) cluster, the Texas Advanced Computing Center (TACC) at The University of Texas at Austin (http://www.tacc.utexas.edu), and the Extreme Science and Engineering Discovery Environment (XSEDE, which is supported by National Science Foundation grant number ACI-1548562) for providing HPC resources that have contributed to the research results reported within this paper.

---

**Figure S1 Single cell RNA-seq data of pancreatic cancer tissues A.** UMAP projections of cells from PDAC-A, which is the paired scRNA-seq data with the spatial transcriptomics data of PDAC-A sample. **B.** UMAP projections of cells from PDAC-B, which is the paired scRNA-seq data with the spatial transcriptomics data of PDAC-B sample. All cells are labeled and colored according to the provided cell type annotations in the original study.

