## Supplementary figures and images for "DSTG: Deconvoluting Spatial Transcriptomics Data through Graph-based Artificial Intelligence"

### Supplemental Figure

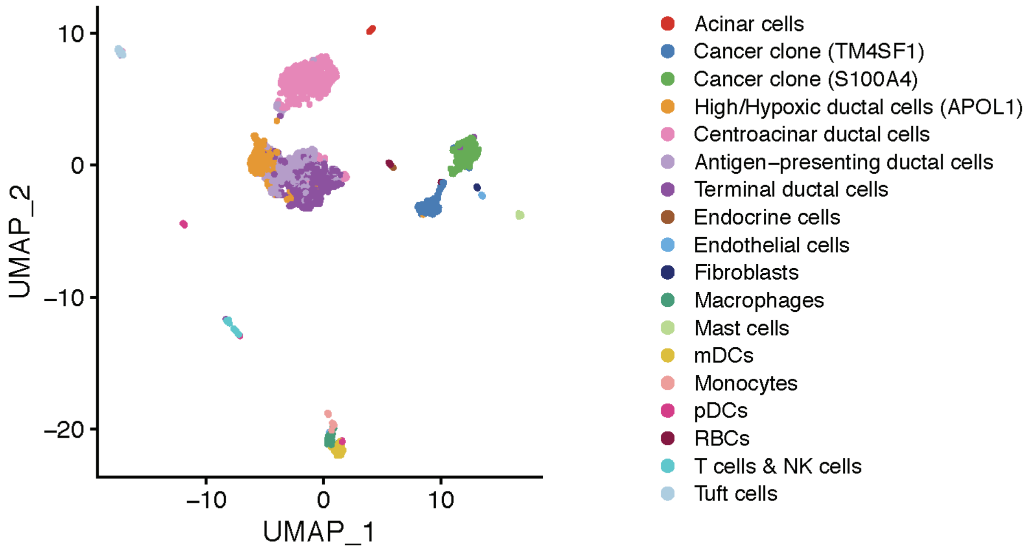

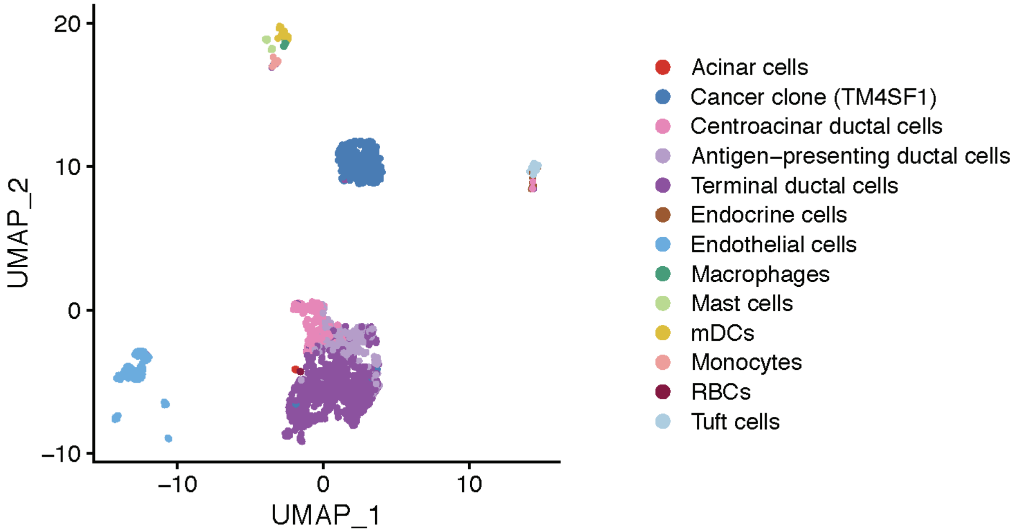
